# The OptoGenBox - a device for long-term optogenetics in *C. elegans*

**DOI:** 10.1101/2020.01.13.903948

**Authors:** Inka Busack, Florian Jordan, Peleg Sapir, Henrik Bringmann

## Abstract

Optogenetics controls neural activity and behavior in living organisms through genetically targetable actuators and light. This method has revolutionized biology and medicine as it allows controlling cells with high temporal and spatial precision. Optogenetics is typically applied only at short time scales, for instance to study specific behaviors. Optogenetically manipulating behavior also gives insights into physiology, as behavior controls systemic physiological processes. For example, arousal and sleep affect aging and health span. To study how behavior controls key physiological processes, behavioral manipulations need to occur at extended time scales. However, methods for long-term optogenetics are scarce and typically require expensive compound microscope setups. Optogenetic experiments can be conducted in many species. Small model animals such as the nematode *C. elegans*, have been instrumental in solving the mechanistic basis of medically important biological processes. We developed OptoGenBox, an affordable stand-alone and simple-to-use device for long-term optogenetic manipulation of *C. elegans*. OptoGenBox provides a controlled environment and is programmable to allow the execution of complex optogenetic manipulations over long experimental times of many days to weeks. To test our device, we investigated how optogenetically increased arousal and optogenetic sleep deprivation affect survival of arrested first larval stage *C. elegans*. We optogenetically activated the nociceptive ASH sensory neurons using ReaChR, thus triggering an escape response and increase in arousal. In addition, we optogenetically inhibited the sleep neuron RIS using ArchT, a condition known to impair sleep. Both optogenetic manipulations reduced survival. Thus, OptoGenBox presents an affordable system to study the long-term consequences of optogenetic manipulations of key biological processes in *C. elegans* and perhaps other small animals.

## Introduction

Optogenetics can control many physiological processes by actively influencing biochemical reactions and manipulating neuronal activity (Fenno, Yizhar, & Deisseroth, 2011). A light-sensitive actuator can be genetically expressed in specific cells of organisms and activated by light. Different tools exist for either activation or inhibition of excitable cells. Some of the most-used tools are channel rhodopsins, which have first been discovered in algae (Nagel et al., 2002, 2003), and ion pumps, which were found in halobacteria (Han et al., 2011), both can now be genetically expressed in other organisms to depolarize or hyperpolarize cells upon light stimulation. Optogenetics has become widely established in different model organisms, e.g. small nematodes and flies but also mammals such as mice and monkeys (Fenno et al., 2011). *C. elegans* is well suited and established for optogenetic studies (Husson, Gottschalk, & Leifer, 2013; Schmitt, Schultheis, Husson, Liewald, & Gottschalk, 2012). Many physiological processes are conserved across species and can be studied in less complex organisms such as the 1mm long nematode *C. elegans*. 83% of its genes have human homologs, allowing molecular studies that are of relevance also to human biology (Lai, Chou, Ch’ang, Liu, & Lin, 2000). With 302 neurons, its nervous system is more manageable than that of other animals. Additionally, a single neuron in *C. elegans* can act similarly to brain regions in mammals (Altun, Z.F. and Hall, 2011). Due to the nematode’s transparency, optogenetic experiments can be conducted in a non-invasive manner (Husson et al., 2013). *C. elegans* was the first animal in which optogenetics was established (Husson et al., 2013; Nagel et al., 2003).

However, there are still limitations that hinder the complete realization of the potential of optogenetics. In particular, long-term optogenetic experiments have rarely been conducted (Schultheis, Liewald, Bamberg, Nagel, & Gottschalk, 2011). In a standard experiment the neuronal manipulation only lasts for seconds or minutes. While it is true that some reactions and neuronal signals are fast acting, to manipulate physiology in the long term, one typically has to manipulate biological processes for days or even longer. Optogenetic long-term experiments are challenging for several reasons:

1. It is necessary to control the environment of the tested organisms.
2. For high-throughput experiments, many different conditions should be processed in parallel.
3. There is currently no inexpensive device available to account for 1 and 2.

Through optogenetic long-term manipulations, it is possible to investigate how a specific behavior affects organisms systemically (Altun, Z.F. and Hall, 2011; Husson et al., 2013; Lai et al., 2000; Schmitt et al., 2012). Even in *C. elegans* research the above-mentioned challenges in long-term optogenetic studies persist. Due to the development of new rhodopsins, first steps towards long-term optogenetics have been made. These newer genetic tools can continually be activated for minutes (Gengyo-Ando et al., 2017) or even for up to 2 days (Schultheis et al., 2011) after a shorter light pulse. The longest optogenetic lifespan experiment to date lasted 2.5 hours (De Rosa et al., 2019).

Optogenetic survival assays lasting several days or weeks have not yet been conducted in *C. elegans*.

One additional reason that explains why long-term experiments have rarely been conducted in *C. elegans* is, that blue light, which is often used in optogenetic

experiments, is harmful to the worms. Blue light causes a negative phototaxis and prolonged exposure leads to paralysis and death of *C. elegans* (Edwards et al., 2008; Ward, Liu, Feng, & Xu, 2008). Alternative optogenetic actuators have been developed that can be excited with a higher wavelength, thus causing less stress to *C. elegans*. For example, the red-shifted Channel Rhodopsin (ReaChR) can be used for neuronal activation (Lin, Knutsen, Muller, Kleinfeld, & Tsien, 2013) or ArchT, which hyperpolarizes neurons by pumping out protons, can be used for inhibition (Okazaki, Sudo, & Takagi, 2012). These genetic tools allow the use of yellow to orange light (585-605nm) for excitation.

Increased arousal and decreased sleep affect the survival of *C. elegans* (De Rosa et al., 2019; Wu, Masurat, Preis, & Bringmann, 2018). Many assays that control arousal and sleep deprivation in *C. elegans* build on external stimuli such as tapping mechanisms, the ablation of neurons or mutation (Bringmann, 2019; Driver, Lamb, Wyner, & Raizen, 2013; Hill, Mansfield, Lopez, Raizen, & Van Buskirk, 2014; Schwarz & Bringmann, 2013; Singh, Ju, Walsh, DiIorio, & Hart, 2014; Spies & Bringmann, 2018; Van Buskirk & Sternberg, 2007). Optogenetics activates or inhibits specific neurons and therefore allows the dissection of neuronal mechanisms. ASH is a nociceptor and its activation causes a reverse escape response by activating the second layer RIM interneurons and by inhibiting the sleep neuron RIS (Kaplan & Horvitz, 1993; Maluck et al., 2020). Mechanical tapping or optogenetic RIM activation, which causes a flight response and increase in arousal, shortens the lifespan of adult *C. elegans* (De Rosa et al., 2019). Depolarization of ASH causes a complex response. It activates RIM, therefore triggering release of tyramine and promoting the flight response (De Rosa et al., 2019; Maluck et al., 2020). Additionally, strong RIM activation inhibits the sleep neuron RIS which leads to sleep deprivation (Maluck et al., 2020). RIS is a single neuron that acts as the motor of sleep in *C. elegans.* RIS is active during sleep, its activation induces sleep and its depolarization is homeostatically regulated (Bringmann, 2018; Maluck et al., 2020; Michal Turek, Lewandrowski, & Bringmann, 2013). A more specific experiment for sleep deprivation, in which arousal also gets increased, is hence the inhibition of the sleep neuron RIS through optogenetics (Maluck et al., 2020; Wu et al., 2018).

To solve the problem of long-term optogenetic manipulation, we have developed the OptoGenBox, a simple-to-use stand-alone device, which provides a controllable environment and allows for the execution of complex optogenetic protocols. The total material costs of less than 3500 USD (Table S1) makes it substantially more inexpensive than the use of standard microscope set-ups. The OptoGenBox therefore presents the currently best solution for long-term optogenetic experiments in *C. elegans*.

We successfully tested the OptoGenBox by optogenetically activating the sensory neuron ASH and inhibiting the sleep neuron RIS. Optogenetic activation of ASH or inhibition of RIS in L1 arrested animals both reduced lifespan. Our results show that the OptoGenBox is a valuable tool for long-term optogenetic experiments in *C. elegans*, and potentially also for other small animals.

## Results

### A device for optogenetic long-term imaging

We developed the OptoGenBox to enable long-term optogenetic experiments in *C. elegans* (Figure 1). Worms were kept in a temperature-controlled environment and illuminated with orange light from the bottom (Figure 2A). For this, the OptoGenBox was built as a 70×70×90cm large device that is programmable via a touch display

**Figure 1.**
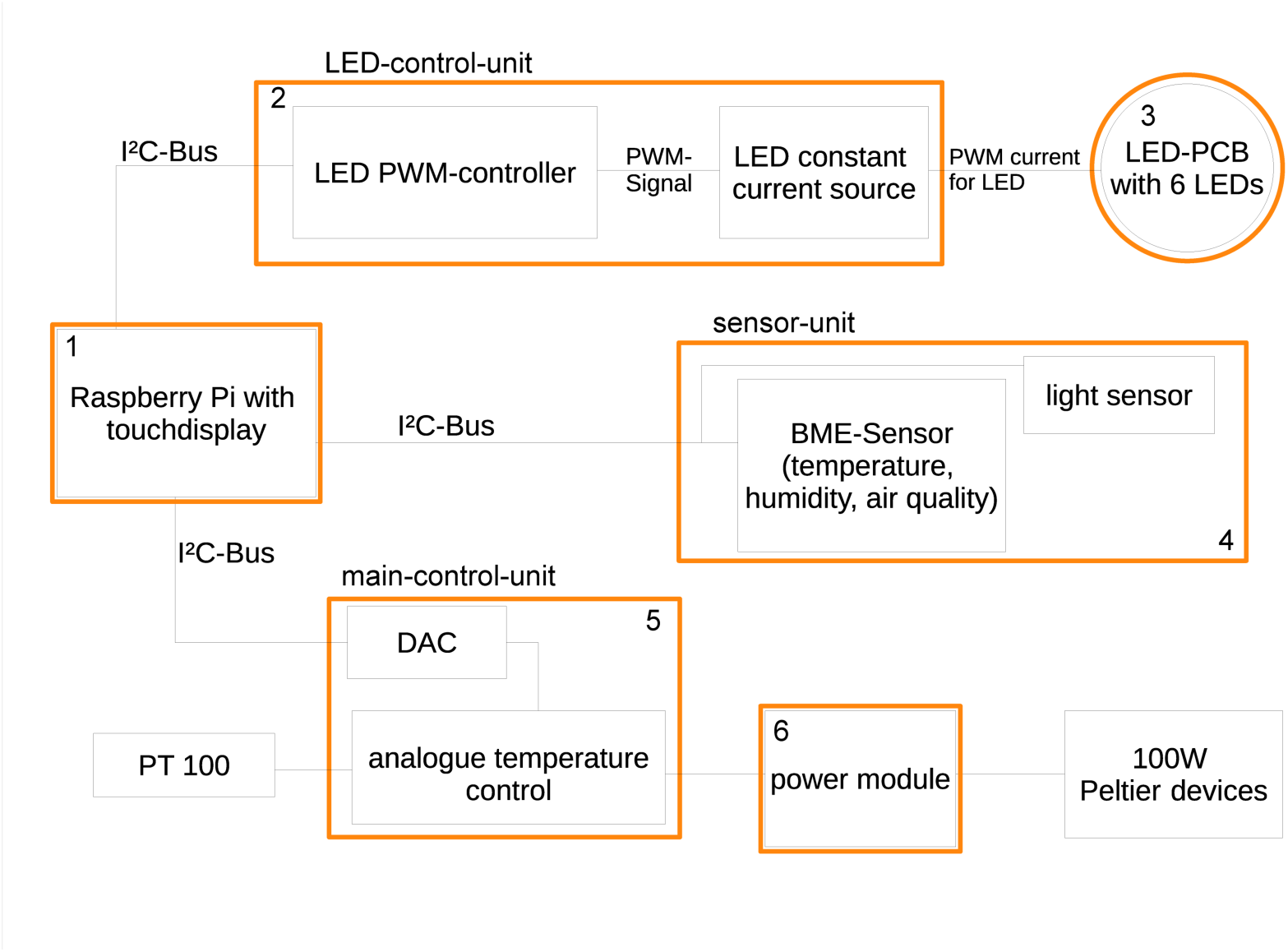
Functional scheme of the OptoGenBox. The box consists of several printed circuit boards (PCBs in coloured outlines), that are connected and controlled by a Raspberry Pi computer.

**Figure 2.**
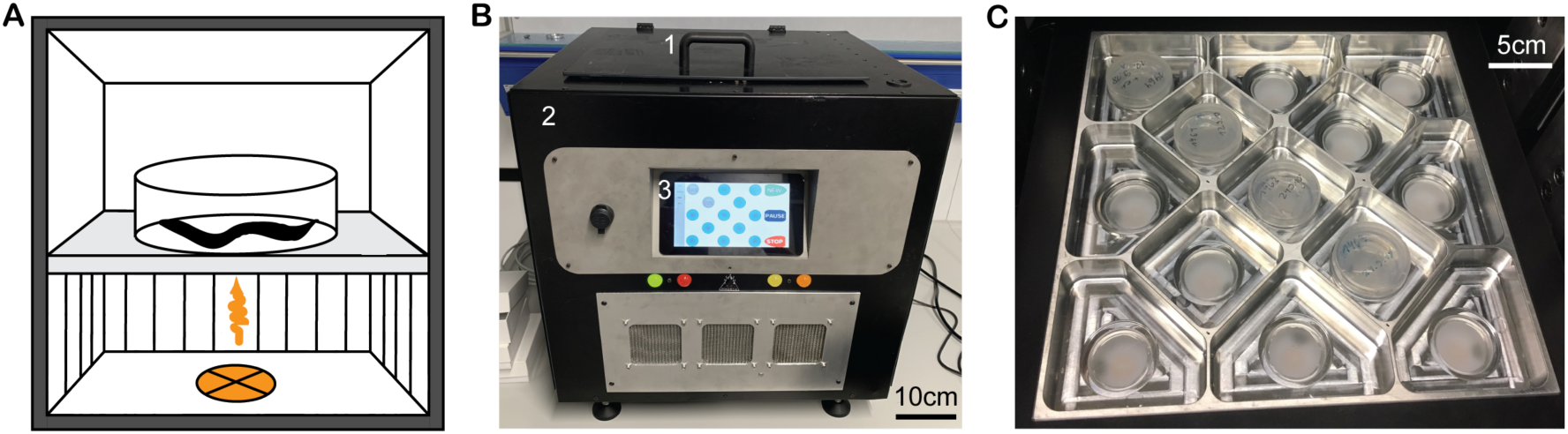
The OptoGenBox is a device for long-term optogenetic experiments in *C. elegans*. A) Worms are kept in a controlled environment and illuminated with orange light. B) The outside of the OptoGenBox. (1) opening handle, (2) exterior case and (3) touch screen. C) The inside of the OptoGenBox is comprised of 13 groupable or separately programmable cells.

(Figure 2B). The inside consists of a 22×22cm sized experimentation area partitioned into 13 cells (Figure 2C). Each cell can hold small plates with nematode growth medium or microfluidic chambers (Bringmann, 2011; M Turek, Besseling, & Bringmann, 2015) with a diameter of 3.5cm, and can thus fit up to 100 worms. Worm plates are placed on 4mm thick glass (B270), which was polished on the bottom (400 polish) to homogeneously distribute the LED light throughout the worm plate (Figure 3). 6 LEDs are distributed throughout an LED module (Figure 4) 7.4mm below the glass to illuminate the worms from the bottom. An aluminum casing keeps external light out and creates optically isolated cells. Furthermore, the box is temperature controlled through Peltier devices and protected from external disturbances via foam and an acrylic case (Figure S1). The LED intensities of all 13 cells were measured with a light voltmeter (ThorLabs PM100A) and calibrated through the software while setting up the system to assure equal light intensities between the cells. The temperature for all cells is uniform and can only be determined when no experiments are running. Each cell contains environmental sensors for light intensity and air quality, and temperature recordings are carried out for each cell. Humidity stays constant in the closed plastic dishes that contain the microfluidic devices (M Turek et al., 2015) and thus humidity measurements are not necessary when the microfluidic devices are used. Nevertheless, sensors are included to monitor humidity inside the device in case other types of samples need to be used.

**Figure 3.**
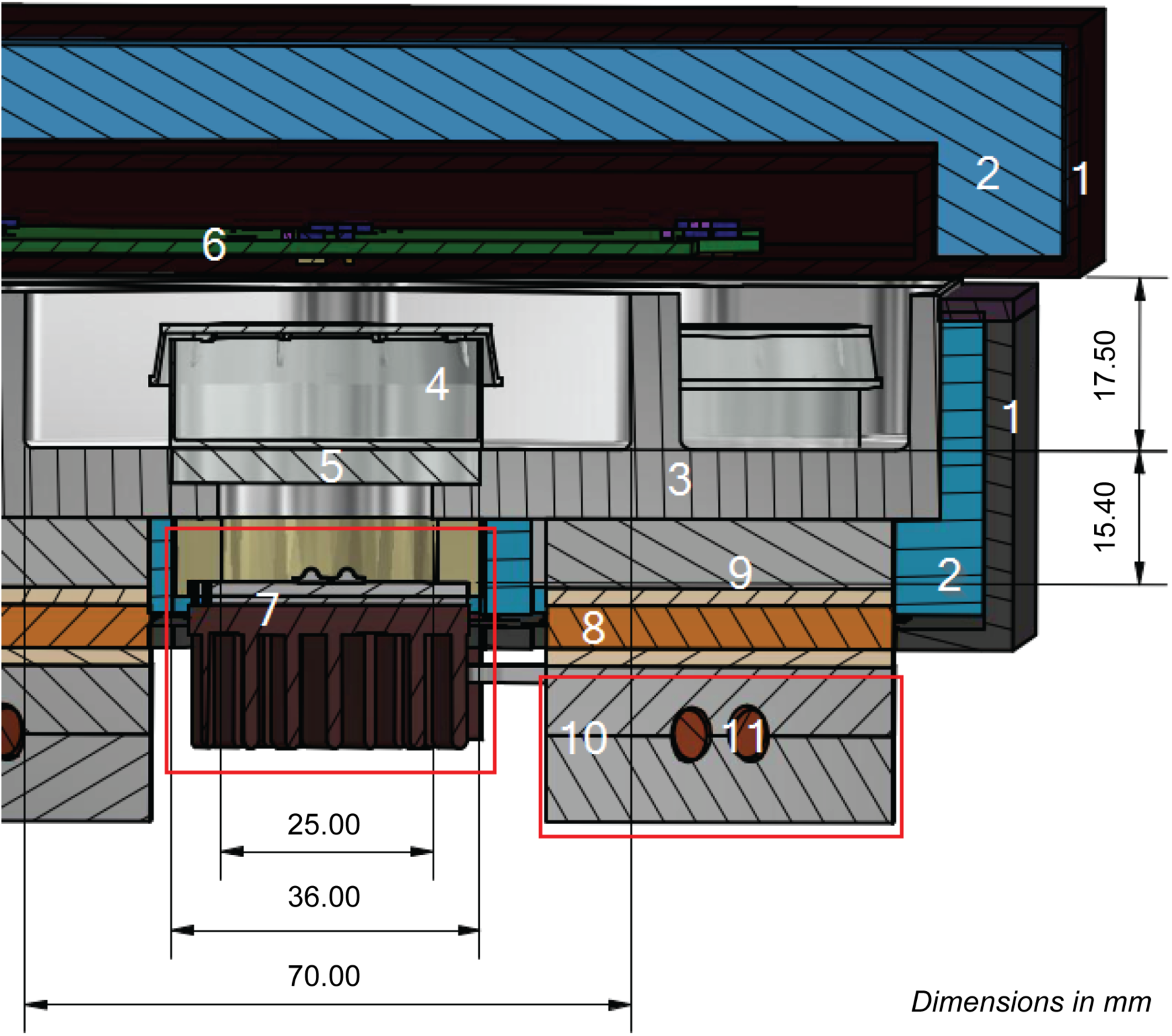
Cross-section of the OptoGenBox incubator. (1) plastic cover, (2) insulating foam, (3) incubator inlet (4) worm plates or microfluidic devices, (5) sanded glass, (6) sensor PCB (in the lid), (7) LED module, (8) mounting bracket Peltier device, (9) Peltier device covered with thermal pads, (10) heat pipe brackets and (11) heat pipe.

**Figure 4.**
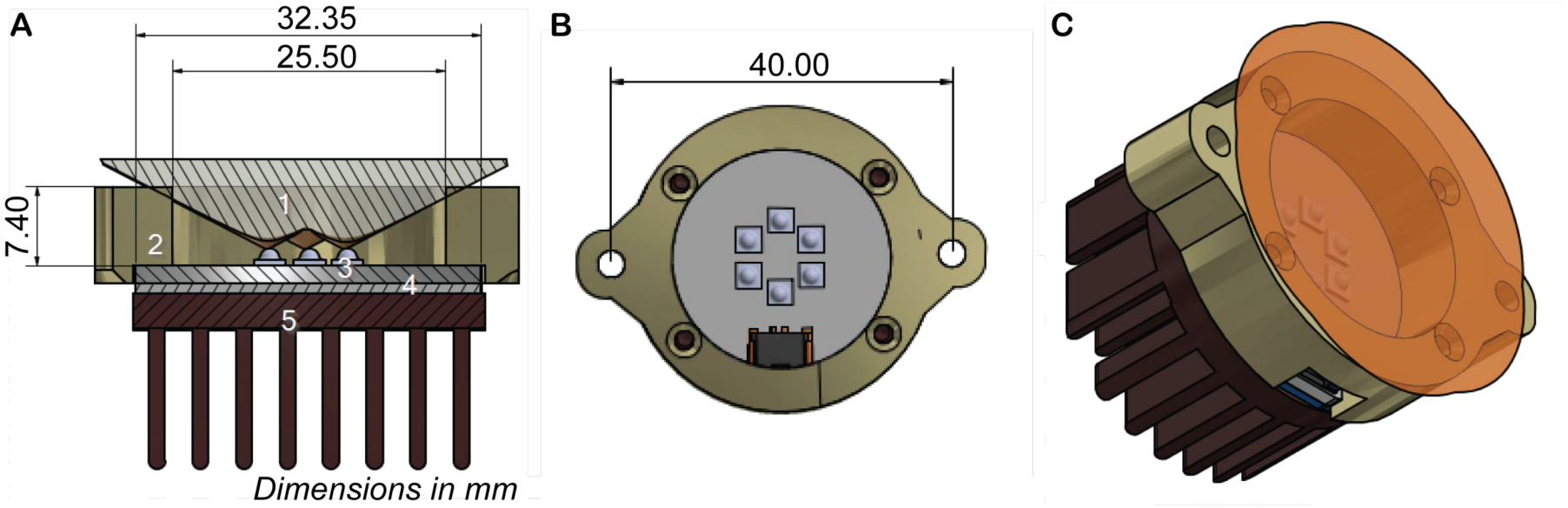
LED module. A) Cross section of an LED module. (1) calculated light beam (LED current at 50%), (2) distance ring to mount the module, (3) the IMS-PCB with single LEDs on it, (4) the thermal pad and (5) the heat sink. B) Top view of the LED module. C) Side view of the LED module with the light distribution.

The researcher can easily program the optogenetic protocol through the touch screen. The system is written in Python and implemented on a Raspberry Pi (Figure S2). To start an experiment the exact cells can be selected individually for each experiment and then the optogenetic protocol can be defined (Figure S3). The experimenter can choose how many cycles should run with how much time (hours or minutes) in light and how much time in darkness and can define the light intensity for the experimentation area (between 2-40mW) during the light times. LEDs can be programmed to be either on or off for minutes or hours. The minimum continuous amount of time for a light cycle is hence 1min and the maximum is 25h. The same holds for dark phases. The maximum number of cycles is 5000. Theoretically, worms could get illuminated for up to 2083 days.

The temperature can be chosen for the entire OptoGenBox between 15-25°C (Figure S4). While one experiment can include up to 13 cells, individual or groups of cells can also be programmed separately to allow for parallel experiments (Figure S5). The total material costs of less than 3500 USD (Table S1) make it much less expensive than microscopic set-ups, which one could also use for optogenetic long-term experiments. All code is freely available (https://gitlab.gwdg.de/psapir/inkubator). The OptoGenbox presents an inexpensive and user-friendly tool to conduct optogenetic long-term experiments.

### Optogenetic ASH activation in OptoGenBox triggers an escape response

The sensory neuron ASH is known to promote reverse escape locomotion upon different harmful stimuli (Kaplan & Horvitz, 1993; Zheng, Brockie, Mellem, Madsen, & Maricq, 1999). To test for the functionality of the box, we developed an escape essay in which we optogenetically activated ASH and tested for its effects on behavior.

ReaChR was genetically expressed in worms under the *sra-6* promoter to cause ASH activation upon addition of ATR (Wu et al., 2018). For the experiment, a small plate was prepared with a small lawn of bacteria of the *E. coli* strain OP50 as a food source on one half of the plate and an opaque sticky tape, which caused an area of shade in the OptoGenBox, on the other half (Figure 5A). Worms without any optogenetic activation were expected to mostly assemble by the food. On the contrary, after ASH activation worms were expected to not gather at the food but to either distribute throughout the plate or gather in the shade, where the activation is interrupted. An optogenetic protocol was run for one hour and the distribution of worms was counted. Indeed, an average of 80% of the control worms gathered by the food. Only around 20% of the ASH-activated *C. elegans* could be counted at the food drop. This significant decrease in worms at the food drop confirms that the worms show an escape response upon ASH activation. Worms did not aggregate in the shade caused by the sticky tape but mostly distributed across the plate. This could potentially be explained by a remaining low light intensity of 0.02mW in the shade (outside the shade there was an intensity of 10mW, so 0.2% of the light intensity could be measured above the sticky tape), which may have still been sufficient for ASH activation and hence an escape response of the worm. The low light intensity in the shade could perhaps be caused by light reflections. Neither the worms in which ASH was activated nor control worms were able to flee from the plate (Figure 5B). These results demonstrate the functionality of the OptoGenBox.

**Figure 5.**
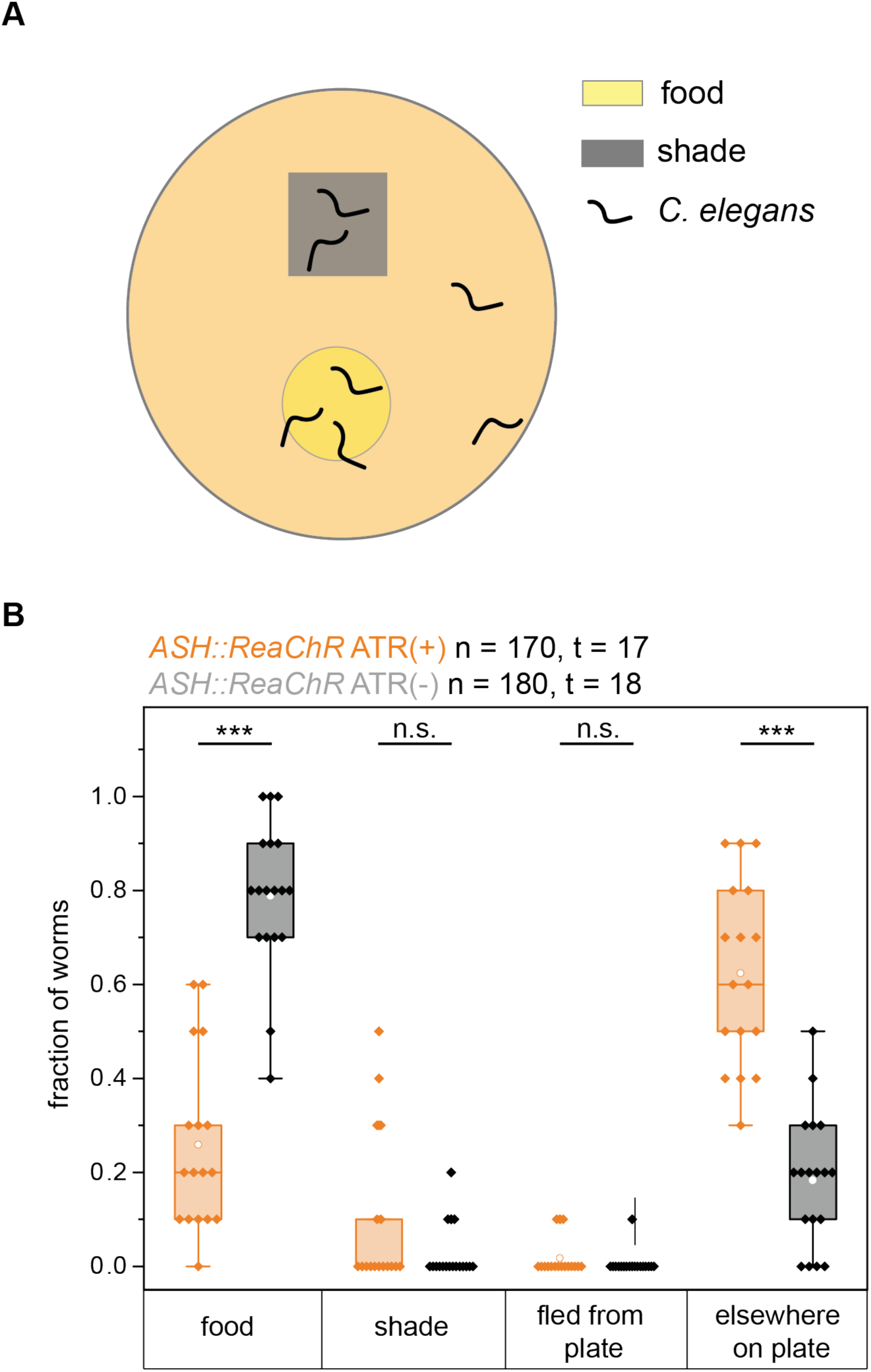
ASH activation in the OptoGenBox caused an escape response. A) Preparation of the experimental plate. A small NGM plate is prepared with drop of food (*E. coli* OP50). An opaque sticky tape is used to block the stimulating light. B) After ASH activation through ReaChR and ATR, worms did not stay on the food drop but distributed throughout the plate. Neither ASH-activated nor control animals fled from the plate. ***p<0.001, Kolmogorov Smirnov Test.

### Increased arousal and decreased sleep by optogenetic manipulations shortens the lifespan of arrested L1 larvae

Increased arousal and sleep deprivation has been shown to shorten the lifespan in *C. elegans* (De Rosa et al., 2019; Wu et al., 2018). We wanted to test if an increase in arousal or inhibition of sleep can affect the survival of arrested L1 larvae. We therefore conducted experiments in which arousal gets increased or sleep is reduced through different optogenetic manipulations.

The optogenetic manipulations were achieved by treating transgenic worms carrying the optogenetic tool with ATR. Since a toxicity of ATR could not be excluded we first investigated the effects of ATR on the wild type. Two rounds of experiments confirmed that the addition of ATR without optogenetic manipulation did not lead to a

significant reduction of survival (Figure 6A and S7A). Hence, any lifespan phenotypes in our optogenetic experiments can be attributed to the optogenetic manipulations and not the treatment with ATR.

**Figure 6.**
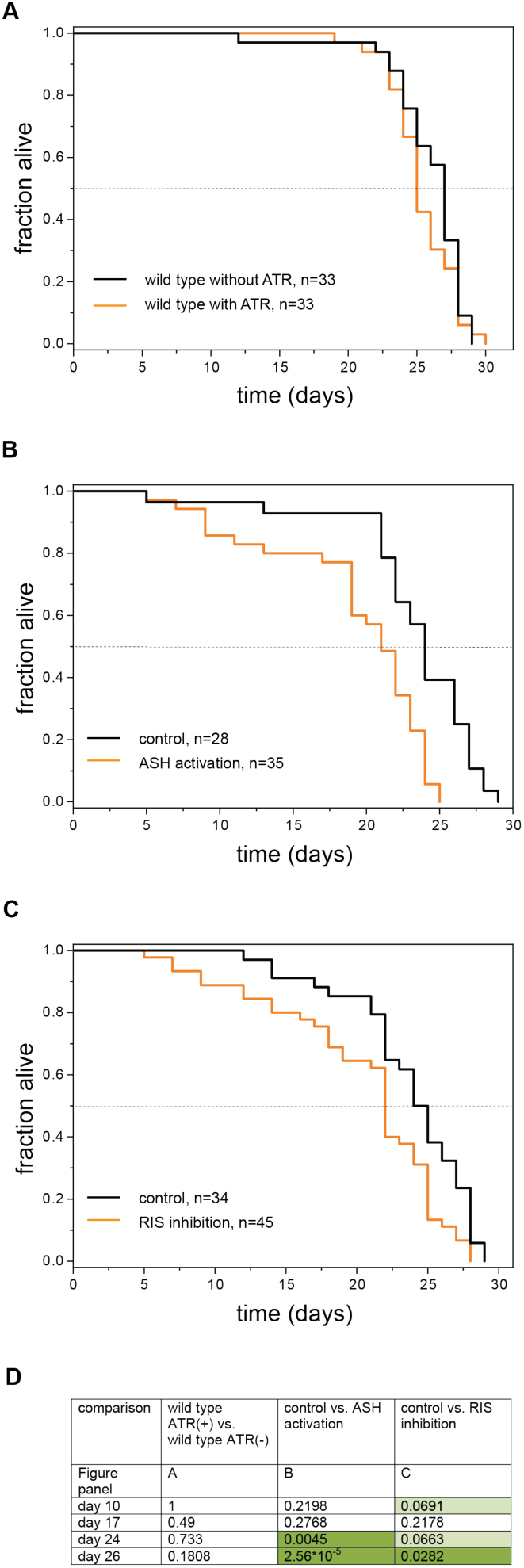
An increase in arousal and sleep deprivation reduces the lifespan of arrested L1 *C. elegans*. A) All-trans retinal (ATR) did not affect survival of wild-type arrested L1 larvae. B) ASH activation causes a reduction in lifespan compared to control animals without the addition of ATR. C) RIS inhibition causes a reduction in lifespan compared to control animals without the addition of ATR. D) p-values of a statistical analysis of lifespans from A-C. Fisher’s Exact Test was conducted at different time points. The p-value of dark green shaded time points is below 0.05 and therefore statistically significant. The p-value of light green shaded time points is above 0.05 but below 0.07 and hence not statistically significant but very close to significance. ASH activation and RIS inhibition caused a significantly decreased lifespan in the later phases of the lifespan.

To test for survival phenotypes upon increased arousal, we conducted two experiments, a first experiment in which optogenetic activation of a nociceptive neuron causes an escape response and increases arousal and a second experiment in which optogenetic inhibition of a sleep neuron causes sleep deprivation.

For the optogenetic activation experiment, we used the ASH::ReaChR strain as described before (Wu et al., 2018). All-trans retinal (ATR) was present throughout the L1 arrest lifespan to ensure functionality of the optogenetic tool. Control worms were used that carried the ReaChR transgene but did not receive ATR. In both rounds of the experiment, animals in which ASH was activated died significantly earlier than control worms (Figure 6B, Figure S7B).

Next, we tested how sleep deprivation caused by the inhibition of the sleep neuron RIS affects survival in L1 arrest. We expressed ArchT under the *flp-11* promoter so that it was specifically expressed in RIS and all-trans retinal was supplemented (Wu et al., 2018). Again, control worms for comparison did not receive ATR treatment. Optogenetic sleep deprivation led to a small but significant reduction of survival in arrested L1 animals by 8.3 % (Figure 6C and S7C).

## Discussion

Here we developed the OptoGenBox as a device for optogenetic long-term experiments. The OptoGenBox combines a controlled environment and allows for parallel processing of many experiments for *C. elegans* and perhaps other small animal models. With material costs of less than 3500 USD it is rather inexpensive. While there exist lower cost alternatives such as the DART system for *Drosophila* (Faville, Kottler, Goodhill, Shaw, & Van Swinderen, 2015), the DART system does not allow for parallel processing and temperature control. Hence, the OptoGenBox currently presents the best solution to allow for parallel optogenetic long-term experiments. Experiments in the OptoGenBox can last up to several weeks. The device allows for optogenetic long-term experiments in a highly controlled environment. OptoGenBox is not equipped with an imaging system. For performing measurements on the worms, the samples containing the worms thus need to be taken out of the system, which could perturb the measurements. However, an imaging system could be added to the device in the future.

We could demonstrate that different optogenetic manipulations that increase arousal or inhibit sleep have a detrimental effect on *C. elegans*. The activation of the nociceptor ASH led to a reduced survival in L1 arrest. While ASH activation also leads to an inhibition of the sleep neuron RIS (Maluck et al., 2020), the lifespan shortage of ASH activated animals cannot solely be accounted for by sleep inhibition(Maluck et al., 2020). More likely, the reduced survival upon ASH activation is caused by the inhibition of cytoprotective mechanisms through the activation of RIM and release of tyramine as has been previously described (De Rosa et al., 2019).

The optogenetic inhibition of RIS presents a very specific and therefore suitable experiment to investigate the effects of sleep deprivation on *C. elegans*. The shortened survival upon RIS inhibition confirms that sleep plays an essential role in arrested L1 worms as has been previously demonstrated with *aptf-1(gk794)* mutants in which RIS is not functional and with worms in which RIS was genetically ablated (Wu et al., 2018). However, the previously reported phenotypes with *aptf-1(gk794)* mutants were stronger, having a reduction of lifespan of approximately 40% compared to the wild type. In comparison, in the optogenetic lifespans, the reduction of lifespan was rather small (around 8.3%). There might be several reasons for these differences. The previously reported stronger lifespan effects were obtained in liquid cultures whereas during the optogenetic experiments, worms were kept isolated in microfluidic devices, making a direct comparison impossible. Furthermore, genetic sleep deprivation by a loss of functional APTF-1 can be presumed to lead to more severe effects than temporally-restricted optogenetic sleep deprivation. The advantages of optogenetics are that behavior can be controlled with temporal precision. Instead of completely depriving the worms of sleep it is possible to study the effects of periodic sleep deprivation. For the results presented here, a long light phase (11h) was followed by only a short dark phase (1h) in each cycle throughout the lifespan. This is a rather long optogenetic stimulation phase, in which neurons could perhaps get desensitized as desensitization has been shown before in optogenetic experiments in *C. elegans* (Bergs et al., 2018; Berndt, Yizhar, Gunaydin, Hegemann, & Deisseroth, 2009). It is possible that shorter intervals of light/dark phases might be more effective for optogenetic sleep deprivation in future experiments.

In experiments with worms in which RIS function was impaired, it was shown that sleep counteracts aging phenotypes (Wu et al., 2018). It would be interesting to see how aging phenotypes progress when RIS is inhibited optogenetically. Additionally, how exactly sleep counteracts aging and causes premature death needs further investigation.

## Conclusions

With the newly developed OptoGenBox, we have mostly investigated how an increase in arousal and a loss of sleep affects survival in L1 arrest. However, many other questions could be answered with our device. Optogenetics is a method that cannot only be utilized for depolarizing or hyperpolarizing neurons but also any other type of cell such as epidermal or muscle cells. Silencing of body wall muscles for example leads to an inhibition of feeding (Takahashi & Takagi, 2017) and photoablation of epidermal cells causes paralysis in *C. elegans* (Xu & Chisholm, 2016). The OptoGenBox should allow for many optogenetic long-term experiments in *C. elegans* and potentially also other small animals. Long-term optogenetics should thus help understand how behavior affects systemic physiology in the long term.

## Methods

### Development of the OptoGenBox

The OptoGenBox consists of several parts to allow for orange light illumination and temperature control (Figure 1–3). The user can select the cells and set the exact light level through the touch display of the raspberry pi computer. Signals from and to the raspberry pi are transferred by an inter-integrated circuit bus (I2C bus). There are four LED controllers that address the LEDs of cells, which the researcher previously chose. The LED controllers convert the set illumination level into a pulse width modulated (PWM) signal. This signal allows a constant current through the LEDs and their current source so that the selected cell gets illuminated. The PWM current finally supplies 6 single high brightness LEDs on one single LED-PCB. A light sensor for each cell gives feedback to the raspberry pi about the activation and wavelengths of the LEDs.

A digital to analogue converter (DAC) connects the digital temperature signal, set by the user, with the analogue temperature control unit. This unit gets the actual value from a PT 100 temperature sensor located at the bottom of the chamber and regulates the power output for six 100W Peltier devices. With this closed control loop the OptoGenBox can operate at a constant temperature between 15°C-25°C

An additional temperature measuring takes part by several evenly placed environmental sensors located in the lid. These sensors measure temperature, air quality (based on gas measurements, 0-50 is excellent air) and humidity. The obtained temperature is displayed on the screen.

The system is built on several printed circuit boards (PCBs), which are separated by function. These are the LED-controlling, the environment measuring, the analogue temperature control, an analogue power module and an overall supplying PCB.

### Assembly of the OptoGenBox

The OptoGenBox consists of a few electronic units (Figure 1), which are: 1) the raspberry pi inclusive the touch display, 2) LED-control-units, 3) LED modules, 4) a sensor-unit, 5) a main-control-unit and 6) a DC/DC-power-supply-unit, These units were specifically produced for the OptoGenBox (except the raspberry pi with its display). Furthermore, all PCBs were assembled manually at the Max Planck Institute for Biophysical Chemistry (MPI-bpC). The bare PCBs were produced by different distributors available in Germany (market compliant).

To assemble a PCB, a soldering iron was sufficient for most PCBs. However, for some PCBs, a reflow-oven was used either because it was required or for a more reliable and time efficient soldering procedure.

#### Reflow Soldering

Reflow soldering requires a special set of tools, which consists of a disposing tool for the soldering paste, a placing machine (not necessary, but facilitates the procedure), and an oven that heats up to at least 270°C.

#### Hand soldering

Hand soldering doesn’t require as specific tools as reflow soldering but requires more skills from the executing person. To produce reliable PCBs, different types of soldering tips are recommended and a set of tweezers should be available.

After PCB assembly, the PCBs were connected. For different types of signals, different connectors and cables were selected. Every connector has its special crimping tool so that in total four crimping pliers were used. Additionally, a set of screwdrivers and pliers should be available. A digital multi-meter was utilized to adjust the LED voltage and to tune the analogue temperature control circuit.

#### Mechanical assembly

The components of the OptoGenBox were placed in a modified case originally build for a water-cooled PC system (Figure S8). In the lower tier, all of the AC/DC power supplies and the temperature-control-unit are placed. The incubator sits in the upper tier of the case. The main-control-unit and the raspberry-pi are placed around the incubator (Figure 2).

The incubator itself is assembled in the following manner:

The outside of the incubator is a plastic cover (Figure 3, number 1) around an insulating foam material (Figure 3, number 2). These two materials provide for a stable temperature environment in the incubator inlet (Figure 3, number 3). The worm plates or microfluidic devices (Figure 3, number 4) can be placed on a one-side sanded glass (Figure 3, number 5) in the inlet. The lid of the incubator contains the sensor-unit (Figure 3, number 6). The LED modules (Figure 3, number 7) and the Peltier devices (Figure 3, number 8) are placed in cut-outs beneath the inlet and each mounted with two screws. This construction makes it possible to change the pre-assembled LED modules. While exchanging the LEDs, one has to pay attention to match the current and voltage to the new LED type for ideal light results. Matching the electrical parameters can be done via already implemented options on the LED-control-unit and DC/DC-power-supply-units.

For an optimized thermal solution, the Peltier devices are clamped with thermal pads between two brackets. One bracket (Figure 3, number 9) is directly attached to the inlet. The other is a two-piece bracket (Figure 3, number 10) clamping the Peltier devices and holding the heat pipes (Figure 3, number 11). The heat pipes transport the emerging heat when the device is cooling the incubator. The elements holding the heat pipes can be assembled separately. The heat pipes were manually bend from a straight pipe to fit in the shape that was needed. All bracket parts were specifically designed for the OptoGenBox. The LED control units were attached to the plastic cover (with standard bolts and screws) and then wired with the 13 LED modules.

With the LED modules and the Peltier devices attached to the insulated, covered aluminium inlet, it was installed on fitting brackets in the upper tier of the modified PC case. The LED-control-unit was wired to the main control unit and to the power supply for the LEDs at the DC/DC power supply unit

Fans were installed on both sides of the case to avoid a cushion of heat beneath the incubator and to create a constant airflow so the LED modules, heatsinks for Peltier devices and the electronics would not get damaged by elevated temperature.

#### The LED module

One LED module consists of six high-power LEDs (Osram Opto Semiconductors LCY-CLBP Series) with a peak wavelength at approximately 590nm with 80lm each (Figure 4). In order to reach a maximum light power of 40mW we placed 6 LEDs in a circle with a diameter of 8.4mm. The individual LED modules were calibrated after the installation to have the same light intensities. At 10mW, the light intensity difference between the center and the periphery of the experimentation area was measured to be 0.04mW (0.4% difference). The PCB of the LED module is an IMS-Core PCB, (insulated metal substrate) to absorb most of the thermal energy and conduct it through a thermal pad to the attached round heat sink away from the temperature-controlled area.

#### The Lid

The lid is made of insulating foam material covered with plastic. To locate the necessary sensors at the designated position, the lid got a fitting cut-out. In this cut-out, an overall covering PCB with a pair of sensors (light & environment) for each individual chamber was placed. It is directly attached to the plastic that covers the aluminium inlet from above, aligned to small holes so that the light can be detected and measured. Through a separated hole the air-quality is measured. This PCB and the plastic, on which it is mounted, could be modified to add several other functions as for example an IR-camera with an integrated light source.

### C. elegans maintenance

Worms were grown at 20°C on Nematode Growth Medium (NGM) plates. The plates were seeded with *E. coli* OP50, which served as food for the worms (Brenner, 1974). The following strains were used for this study: HBR974 *goeIs232(psra-6::ReaChr::mKate2-unc-54-3’utr, unc-119(+))* HBR1463 *goeIs307(pflp-11::ArchT::SL2mKate2-unc-54-3’utr, unc-119(+))* N2 wild type (Bristol) (Brenner, 1974)

### Escape Assay

Late L4 stage worms were picked onto NGM plates with 0.2 mM all-*trans* retinal (ATR, Sigma Aldrich). Control late L4 stage worms were picked onto NGM plates without ATR. 9.6 cm2 large NGM plates were prepared for the experiment by placing a 1 cm2 opaque tape on the bottom of one side of the plate and a drop of *E. coli* OP50 on the other side (Figure 5A). After 4 hours, 10 young adult worms were picked into the food drop of the experimental plate for each trial.

The experimental plates were then placed in the OptoGenBox and stimulated with 10mW orange light for 1 hour at 20°C. After one hour the plates were removed from the OptoGenBox and the distribution of worms on the plates was counted.

#### Lifespan assay

It was shown before that sleep is important for the survival of *C. elegans* by counteracting aging phenotypes. However, non-sleeping *aptf-1(gk974)* mutants only have a reduced lifespan when worms starve upon hatching and arrest in the first larval stage (L1 arrest) and not when they are adults (Wu et al., 2018). For this reason, we conducted our experiments with L1 arrested animals.

Worms were kept in microfluidic devices as previously described (Bringmann, 2011; M Turek et al., 2015). A PDMS mold was used as a stamp to cast 110×110×10µm cuboids into a hydrogel. The hydrogel consisted of 3% agarose dissolved in S-Basal (Stiernagle, 2006). Eggs were transferred from a growing plate to a plate without food and then picked into chambers without transferring food. Between 29 and 45 worms were in one microfluidic device housed in individual chambers.

For optogenetic activation or inhibition, chambers were replenished with 10µl of 10mM all-trans-retinal (ATR, Sigma Aldrich) every 3-4 days. Control chambers did not

receive ATR. To avoid fungal contamination, 20µl of 10µg/ml nystatin was pipetted to each chamber 2-4 times throughout the lifespan. Additionally, 20µl of sterile water was added all 2 days until day 15 of the lifespan and then each day to counteract the agarose drying out over time. In the beginning of the lifespan experiment, worms were counted every second day, in the later stages of the survival assay they were counted every day. A worm was counted as dead if it didn’t move for 2 min under stimulation with a blue light LED. This was necessary to distinguish dead from sleeping worms.

The worms were placed in the OptoGenBox and illuminated with 10mW for 11h to attain a long continuous neuronal manipulation. This was followed by 1h of darkness to allow the optogenetic tools to recover without giving too much time to sleep homeostasis processes. This protocol was repeated until all worms were dead. The temperature of the incubator was set to 20°C.

### Statistics

Sample sizes were determined empirically based on previous studies. The researcher was not blinded since the addition of ATR is easily detectable. Conditions in the escape assay were compared with the Kolmogorov Smirnov Test. For the lifespan assays 4 standardized time points were compared. Data of both conditions were compared on these days with Fisher’s Exact Test.

## Acknowledgements

We thank the CGC, which is funded by NIH Office of Research Infrastructure Programs (P40 OD010440), for the N2 strain. The mechanics workshop at the MPI BPC provided us with valuable advice for the design and parts of the OptoGenBox. We would also like to thank Juliane Haase for assisting with laboratory work. This work was funded by the Max Planck Society (Max Planck Research Group), a European Research Council Starting Grant (ID: 637860, SLEEPCONTROL), and the University of Marburg.

## Author Contributions

IB and FJ designed the OptoGenBox. IB designed, performed and analyzed the experiments and wrote the manuscript. FJ built the hardware of the OptoGenBox and contributed to the manuscript. PS programmed the software of the OptoGenBox. HB acquired funding, conceived the project, supervised the work, and edited the manuscript.

## Supplementary Material

There are 8 supplementary figures and 1 supplementary table. The code for the OptoGenbox can be found on github (https://gitlab.gwdg.de/psapir/inkubator). Data for the experiments is published on Mendeley Data (http://dx.doi.org/10.17632/d7wfc9fdbb.2).

