## Supplementary Materials for "The OptoGenBox - a device for long-term optogenetics in *C. elegans*"

### The set-up of the OptoGenBox

#### *Thermal control of cells*

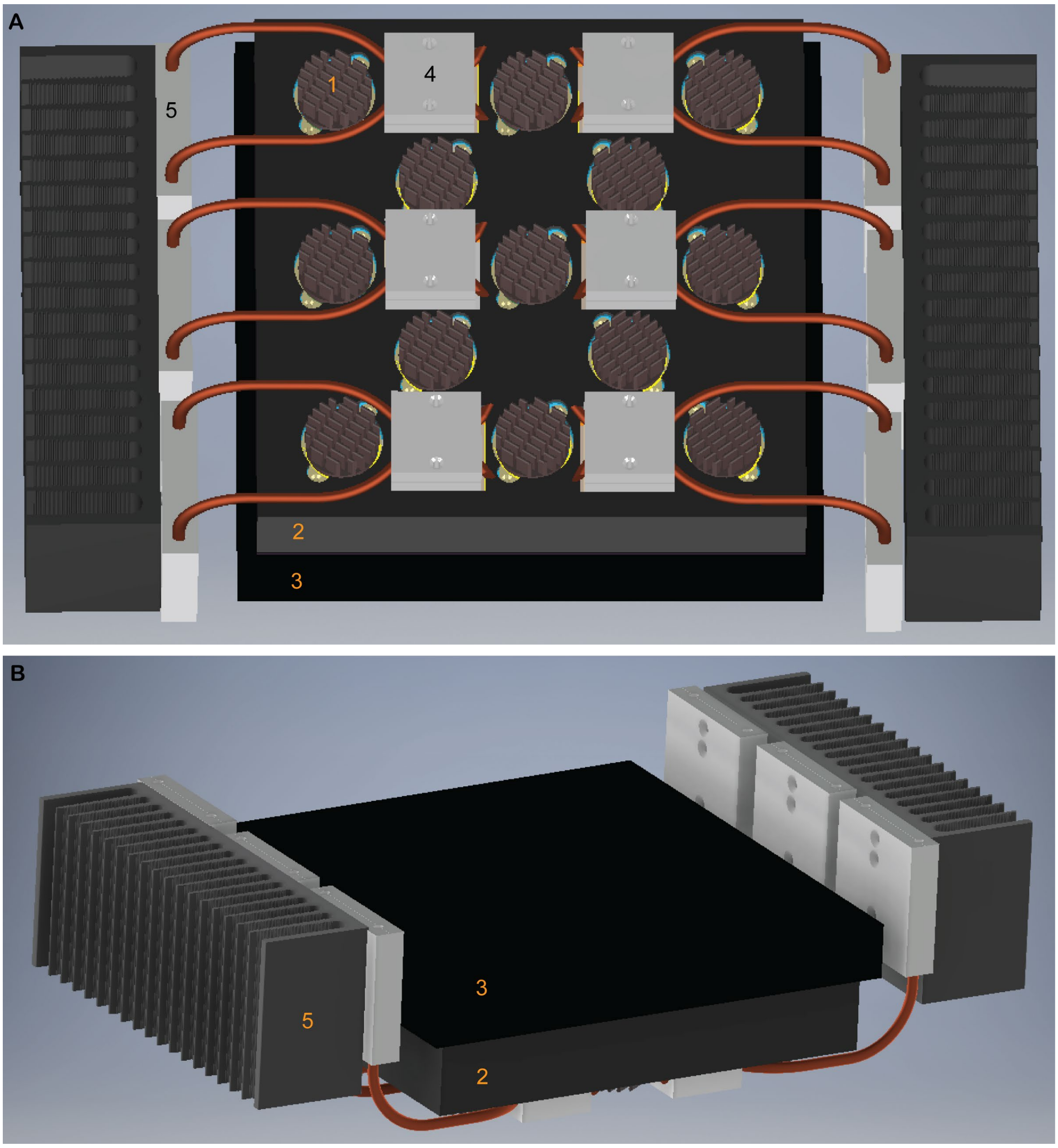


Figure S1. The incubator.

Bottom (A) and top side (B) view of the incubator. The (1) 13 LED modules are spread throughout (2) an incubator chamber and covered by (3) a lid. Six Peltier devices are covered with (4) heat spreaders. On the site are (5) heat sinks.

### Software of the OptoGenBox

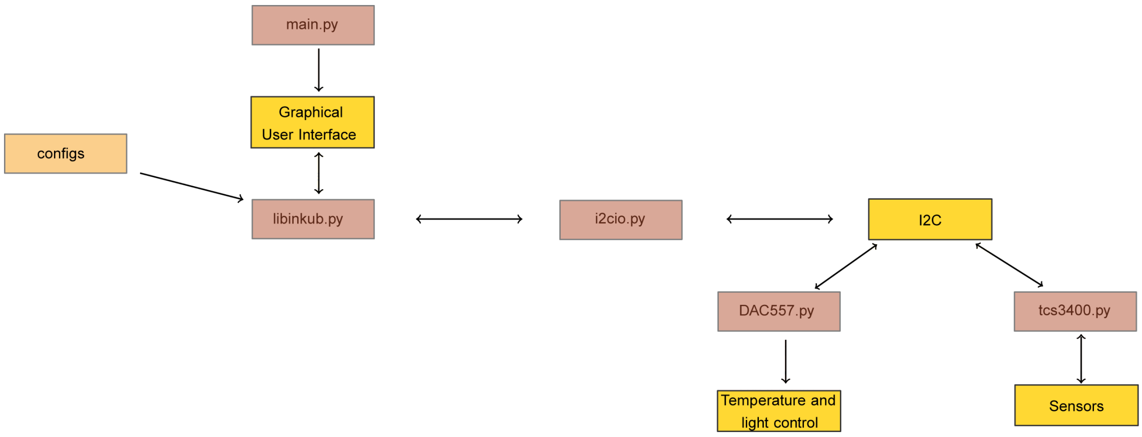


Figure S2. Scheme for the code of the OptoGenBox.

The software was written in python. The python library i2cio reads and writes commands via the device I²C bus. The python library tcs3400 controls the lightsensor TCS3400. The digital-to-analogue converter DAC5571 is needed for the temperature control and controlled by the DAC5571 python library. The main library starts the graphical user interface. The libinkub library manages the abstraction of cells and group of cells, light cycles and sensor readings. The configs file hosts all configurations for the OptoGenBox such as the users, the temperature and light intensity steps and how thoroughly the sensors get calibrated (how many calibration points get initiated). The code is published on github (https://gitlab.gwdg.de/psapir/inkubator).

*Graphical User Interface*

The OptoGenBox program starts by running the main script in the inkub directory in the terminal with the command lines:

```
cd inkub
./main.py
```

The program initiates by calibrating the environmental sensors. This might take a few minutes.

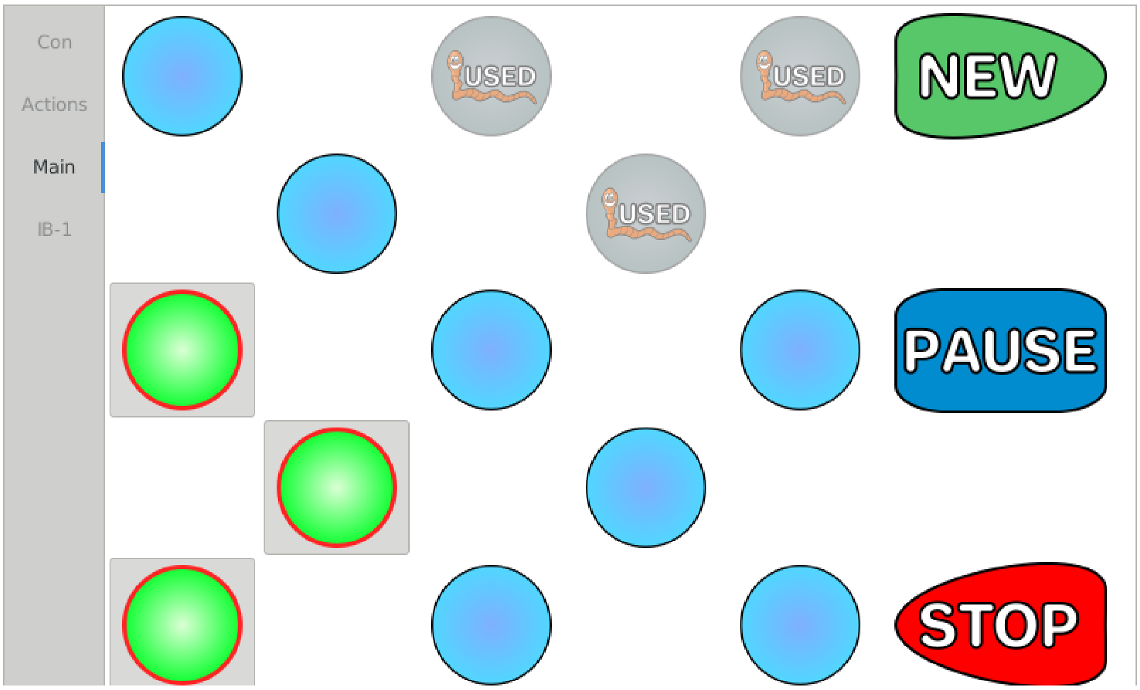


Figure S3. Main menu.

The graphical user interface allows for an intuitive usage of the OptoGenBox. The researcher first selects the cells for one experiment via the touch screen in the main menu and presses *NEW*. Once an optogenetic protocol (light and dark cycles) is defined for selected cells, these cells are displayed as used in the main menu.

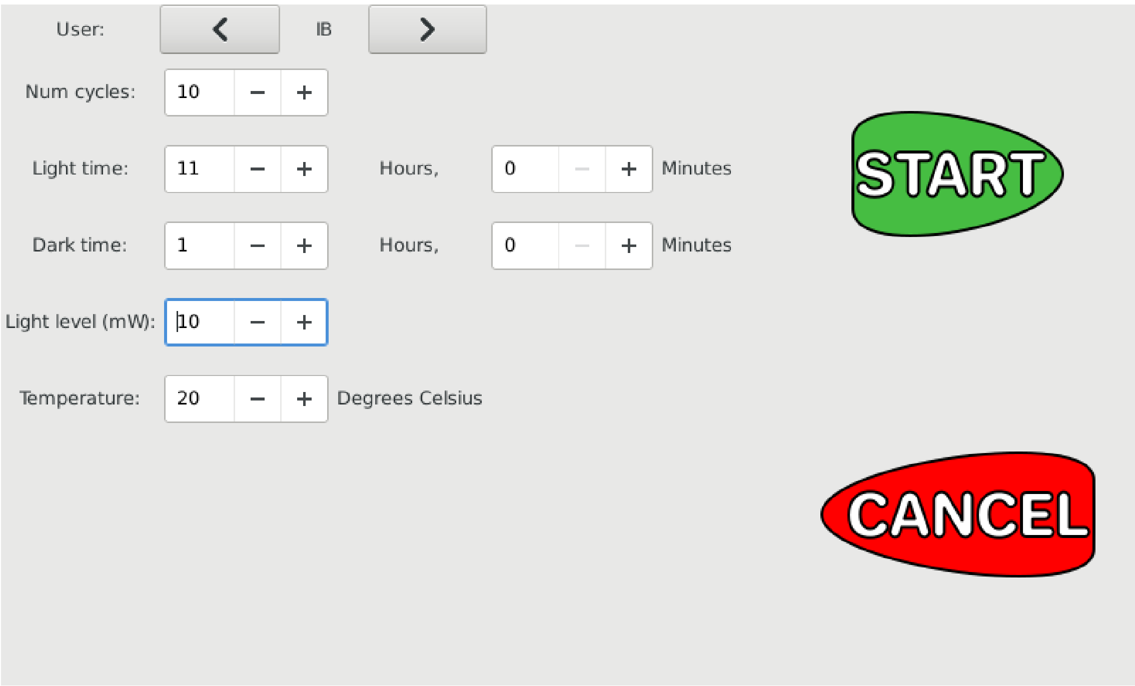


Figure S4. Optogenetic Protocol.

A new window opens and the researcher can select the user and the number of cycles. Furthermore, the researcher can define the time of light and darkness and the level of light. The temperature of the entire incubator can be chosen whenever no other experiments are running. Once all parameters are defined, the protocol gets started by pressing *START* on the touch screen.

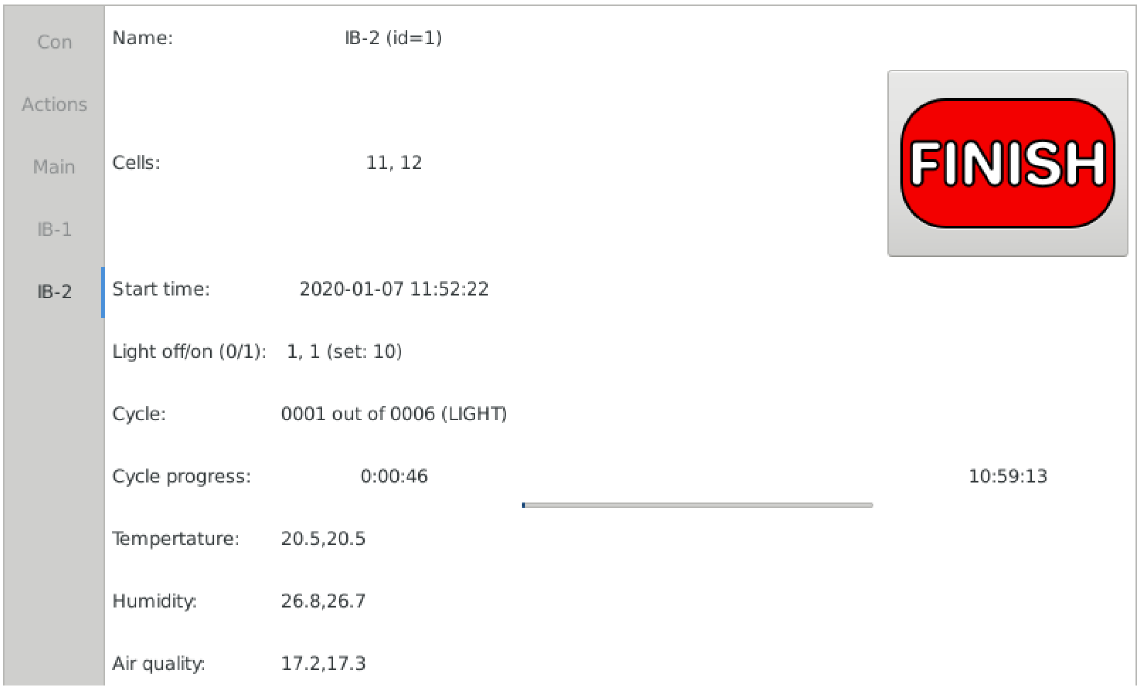


Figure S5. Status of the running protocol.

Running protocols are listed in the left panel below *Main.* To check for the current status of the protocol, one can select it via the touch screen. Information about the name of the experiment, the selected cells and the start time is portrayed. Furthermore, one can see if the light is on or off for each cell and what intensity is set. The next line gives the number of the current cycle. Below that one can see the exact timing in the current cycle. The measured temperature, humidity and air quality is additionally given for each cell. To prematurely quit the experiment one can click the *FINISH* button.

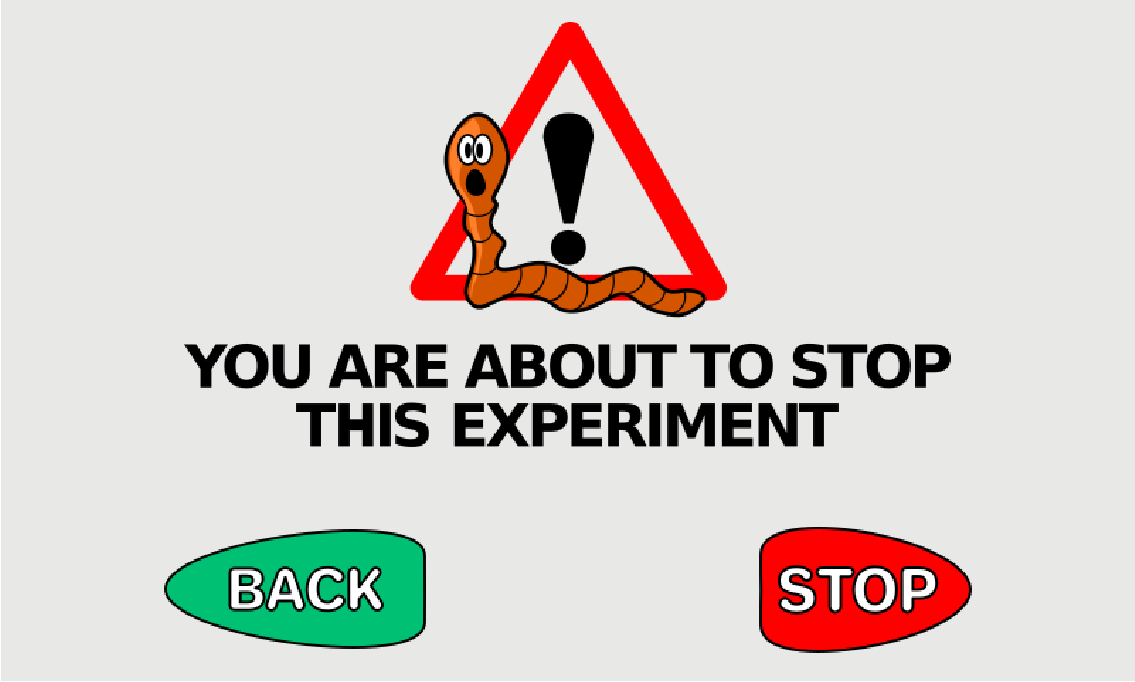


Figure S6. Finishing a running experiment.

Clicking the *FINISH* button on the running protocol screen does not automatically end the experiment. The user has to confirm again that the protocol should be stopped.

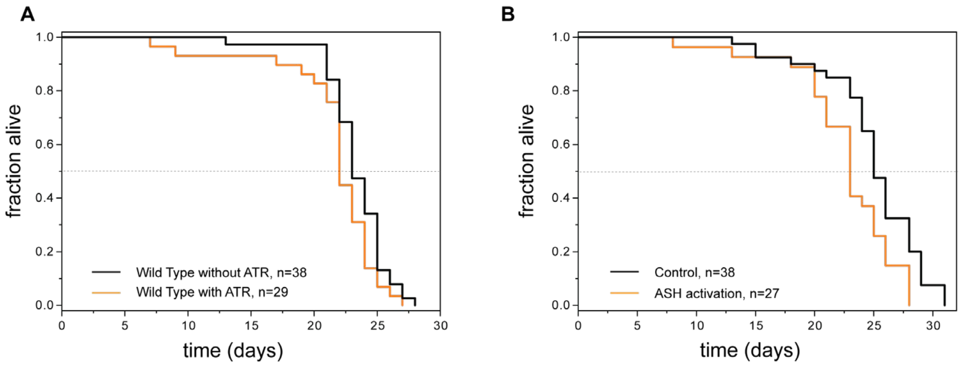


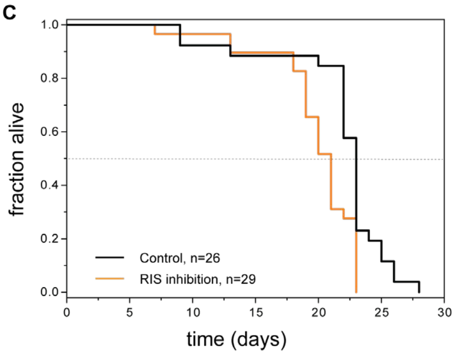


D

| comparison | wild type ATR(+) vs. wild type ATR(-) | control vs. ASH activation | control vs. RIS inhibition |
|---|---|---|---|
| Figure panel | A | B | C |
| day 10 | 0.189 | 0.409 | 0.5908 |
| day 17 | 0.5779 | 1 | 1 |
| day 24 | 0.1335 | 0.0044 | 0.0069 |
| day 26 | 0.453 | 0.0461 | 0.09 |

Figure S7. Second batch of lifespan experiments.

A) All-trans retinal (ATR) did not affect survival of wild-type *C. elegans* in L1 arrest.

B) ASH activation causes a reduction in lifespan. compared to control animals without the addition of ATR.

C) RIS inhibition causes a reduction in lifespan compared to control animals without the addition of ATR.

D) p-values of a statistical analysis of lifespans from A-C. Fisher's Exact Test was conducted at different time points. The p-value of dark green shaded time points is below 0.05 and therefore statistically significant. The p-value of light green shaded time points is above 0.05 but below 0.1 and hence not statistically significant but close to significance. ASH activation and RIS inhibition causes a significantly decreased lifespan in the later phases of the lifespan.

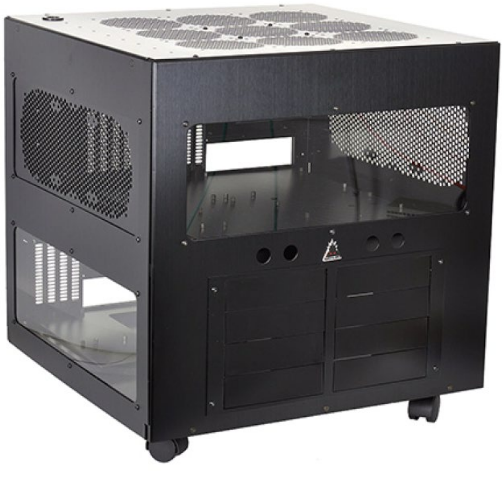

Figure S8. Phobya WaCoolT Cube 2 Watercase was modified to serve as external case for the OptoGenBox. This figure was reproduced with permission from Performance PCs.

| description | manufacturer | manufacturer number | # | price, total $ |
|---|---|---|---|---|
| ***mechanical parts*** | | | | |
| case | Phobya | WaCool IT 2 | 1 | $146 |
| heat pipes | QuickCool | QY-SHP-D6-250SA | 12 | $127 |
| heat sink | Fischer elektronics | SK580 200 SA | 2 | $78 |
| dust filter | InLine | 33378A | 3 | $7 |
| LED-mounting Ring | MPI-bpc ES | inhouse design | 13 | $130 |
| mounting material | | miscellaneous | | $336 |
| material: percision mechanics & optics | | inhouse design | | $538 |
| ***computer elements and accessories*** | | | | |
| raspberry pi | Raspberry Pi | Raspberry Pi 2 Model B | 1 | $22 |
| touchscreen | Raspberry Pi | 7“, 800x400Pixel | 1 | $78 |
| usb connectors | | miscellaneous | 2 | $34 |
| RJ-45 connector | Harting | Harting: 09454521561 | 1 | $17 |
| fan 80mm | Be quiet | BQT BL044 | 5 | $34 |
| fan 120mm | Be quiet | BQT BL046 | 4 | $32 |
| ***power sources*** | | | | |
| Power Adapter AC/DC 330W | Mean Well | HRPG-300-15 | 2 | $203 |
| Power Adapter AC/DC 200W | Mean Well | SP-240-24 | 1 | $63 |
| Power Adapter DC/DC | Traco Power | TEL3-2022 | 1 | $29 |
| Power Apdater DC/DC | Traco Power | TSR3-24150 | 1 | $38 |
| Power Adapter DC/DC | TDK Lambda | I6A-240-14A-033V/001 | 2 | $105 |
| PCB (supply) & small parts | MPI-bpc ES | inhouse design | 1 | $61 |
| ***LED module*** | | | | |
| PCB | MPI-bpc ES | inhouse design | 13 | $131 |
| LED: LCY-CLBP | Osram opto Semicondutcors | LCY-CLBP KXKZ-5F5G | 78 | $131 |
| connectors | ERNI | Erni MiniBridge,2-pin | 13 | $22 |
| heatsink | Fischer elektronics | ICK_LED R33x16,5G | 13 | $121 |
| ***LED-Control-Unit*** | | | | |
| PCB | MPI-bpc ES | inhouse design | 2 | $157 |
| Controller IC [I²C] | NXP Semiconductors | PCA9685 | 4 | $9 |

| | | | | |
|---|---|---|---|---|
| constant current source | ON Semiconductors | CAT4101 | 14 | $39 |
| other electronical parts | | miscellaneous | 158 | $49 |
| ***Heating & cooling*** | | | | |
| Peltier devices 100W | True Components | HP-127120-40x40 | 6 | $134 |
| Power Amplifiers | Texas Instruments | OPA541AP | 3 | $61 |
| PCB | MPI-bpc ES | inhouse design | 1 | $67 |
| temperature sensor | RS Pro | PT100, 8x2mm | 1 | $11 |
| other electronical parts | | miscellaneous | 38 | $56 |
| ***Main control-unit*** | | | | |
| PCB control | MPI-bpc ES | inhouse design | 1 | $78 |
| other electronical parts | | miscellaneous | 85 | $45 |
| ***Lid-&Sensor Unit*** | | | | |
| PCB | MPI-bpc ES | inhouse design | 1 | $146 |
| Light-Sensors | AMS | TCS3400 | 13 | $36 |
| Environment Sensors | Bosch Sensortec | BME680 | 13 | $110 |
| Multiplexer [I²C] | NXP Semiconductors | PCA9548A | 2 | $3 |
| other electronical parts | | miscellaneous | 40 | $11 |
| | | | | **$3,496** |

**Table S1. Material list for the OptoGenBox.**

**Abbreviations**

ATR - all-trans retinal

DAC -  digital-to-analogue converter

$I^2C$ – inter-integrated circuit

IMS – insulated metal substrate

LED – light-emitting diode

NGM- nematode growths medium

PCB – printed circuit board

PWM – pulse width modulation
